# An endolithic *Gloeobacter* and other novel cyanobacterial diversity in alpine dolomites of Piora Valley

**DOI:** 10.64898/2026.10.08.757548

**Authors:** Edi Sudianto, Benoit Durieu, Luc Cornet

## Abstract

**Background:** Cyanobacteria are the major phototrophs in many endolithic ecosystems, yet their genomic diversity and metabolic adaptations remain poorly characterized. Here, we investigated cyanobacterial diversity in endolithic dolomite rocks from the Piora Valley in the southern Swiss Alps using genome-resolved metagenomics.

**Results:** We reconstructed 138 medium-quality metagenome-assembled genomes (MAGs) from four neighboring endolithic rock communities, including 15 cyanobacterial MAGs distributed across six order-level lineages. The cyanobacterial communities showed limited taxonomic overlap among rocks, and most recovered MAGs represented potentially novel species. Only one *Gloeobacter* lineage occurred in all sampled rocks, which we propose as *Gloeobacter dolomitensis* sp. nov., representing the first *Gloeobacter* genome recovered from an endolithic environment. Comparative genomic analyses identified a *cynS* gene encoding cyanase unique to *G. dolomitensis* among the *Gloeobacter*.

Phylogenetic analyses support its horizontal acquisition from Pseudomonadota. Interestingly, the genome of *G. dolomitensis* lacks a canonical CynABD cyanate transporter and instead contains a distinct ATP-synthase binding cassette (ABC) transporter adjacent to *cynS*. Although the available genomic and structural evidence does not establish cyanate transport, its genomic association with *cynS* suggests a potentially non-canonical mechanism. We further show that horizontal gene transfer was unevenly distributed among cyanobacterial lineages, with Acaryochloridales acting predominantly as donors, Chroococcidiopsidales as recipients, and Gloeobacterales showing a comparatively high frequency of interphylum gene exchange.

**Conclusions:** The Piora Valley dolomite environment represents an underexplored reservoir of cyanobacterial diversity, including basal and previously undescribed lineages. The discovery of *G. dolomitensis* and its unusual cyanate-associated genomic architecture expands our understanding of Gloeobacterales diversity and highlights the potential importance of horizontal gene acquisition and metabolic innovation in endolithic microbial communities.

## Background

Endolithic microorganisms live within rocks, occupying protected subsurface niches where the mineral matrix creates a distinct microbial habitat at the interface between terrestrial life and the geosphere [1]. Although life inside rocks is characterized by oligotrophic conditions, desiccation, and intense radiation, the rock matrix can buffer environmental extremes and provide a refuge for microbial life [2]. Endolithic communities have been identified across diverse cold, arid, aquatic, and terrestrial environments worldwide (reviewed in [3]). Cyanobacteria are major phototrophic members of these communities and have therefore been a central focus of endolithic microbiology. Endolithic cyanobacteria have been reported from numerous habitats and include members of *Chroococcidiopsis, Gloeocapsa, Leptolyngbya, Synechocystis*, and *Synechococcus* [4]. Notably, the oldest microorganism identified from an endolithic environment is the cyanobacterium *Eohyella campbelli*, discovered in approximately 1.5-billion-year-old rocks of the Dahongyu Formation in China [5].

One well-characterized endolithic habitat is the Piora Valley in southern Switzerland, where exposed, porous dolomite rocks harbor distinct microbial bands a few millimeters below the rock surface [6]. These bands are dominated by phototrophs and range from grayish-green to red-green in appearance [6]. Molecular and microscopic studies have revealed diverse cyanobacteria in the Piora Valley, including *Leptolyngbya, Nostoc, Scytonema, Microcoleus, Chroococcidiopsis, Anabaena*, and *Gloeobacter* [6–8]. However, these studies relied primarily on microscopy, cultivation, and short 16S rRNA gene sequences, providing limited taxonomic resolution and little information about the functional potential or genomic adaptations of the endolithic community. Thus, despite the well-documented cyanobacterial diversity in Piora Valley dolomite, the genomic diversity and metabolic capabilities underlying persistence in this environment remain poorly understood.

Among cyanobacteria, early-branching lineages are of particular interest for understanding the evolution of oxygenic photosynthesis. Gloeobacterales is one of the basal lineages of extant photosynthetic cyanobacteria and is distinguished by the absence of thylakoid membranes, with photosynthetic complexes localized instead in the cytoplasmic membrane [9,10]. Other early-branching cyanobacterial lineages, including Gloeomargaritales and Thermostichales, provide additional phylogenetic context for understanding the emergence and evolution of oxygenic photosynthesis [11]. Historically, these lineages were represented by few cultured organisms and genomes, limiting our understanding of their diversity and ecology. Recent studies expanded this picture, revealing previously unrecognized Gloeobacterales and other early-branching cyanobacteria from diverse environments, including polar habitats, freshwater systems, soils, caves, and rock-associated environments [12]. For example, Grettenberger [13] recovered five previously unreported Gloeobacterales genomes from Arctic soil, a freshwater spring, peat, and a Hawaiian cave, substantially expanding the genomic representation of this sparsely sampled lineage. More recently, large-scale metagenomic surveys further indicate that early-branching cyanobacteria are predominantly associated with non-marine environments and remain underrepresented in conventional environmental datasets [12]. Together, these findings suggest that the apparent rarity of early-branching cyanobacteria may reflect historical undersampling and limited genomic representation rather than a restricted ecological distribution.

Here, we use genome-resolved metagenomics to characterize the cyanobacterial diversity inhabiting endolithic dolomite rocks in the Piora Valley. We recover 15 cyanobacterial MAGs spanning six order-level lineages, including previously uncharacterized Gloeobacterales genomes, and describe a novel *Gloeobacter* species, *Gloeobacter dolomitensis*. We further identify *G. dolomitensis* as the only *Gloeobacter* genome recovered across all sampled rock communities. The genome of *G. dolomitensis* contains a horizontally acquired *cynS* gene encoding cyanase and is associated with a divergent, potentially non-canonical ATP-synthase binding cassette (ABC) transporter. These findings provide evidence for a previously unrecognized cyanate-associated genomic system in Gloeobacterales and raise the possibility of cyanate utilization or detoxification in this lineage.

## Methods

### Rock sampling, DNA isolation, and sequencing

Dolomite rock samples used in this study were collected from the Piora Valley in the southern Swiss Alps (46°32′51′′ N, 8°43′05′′ E; Figure 1) at an elevation of 1,965 m above sea level in 2025. Rock samples were collected approximately 1 m from the cave entrance. The rocks were mechanically split using a sterile chisel and hammer. Rock fragments displaying clearly visible endolithic bands were selected and transferred into sterile Falcon tubes. Samples were stored in the dark at 5°C until further processing. The sampling map was drawn using ggspatial v1.1.10 [14] and rnaturalearth v1.2.0 [15] in R v4.1.2 [16].

**Figure 1.**
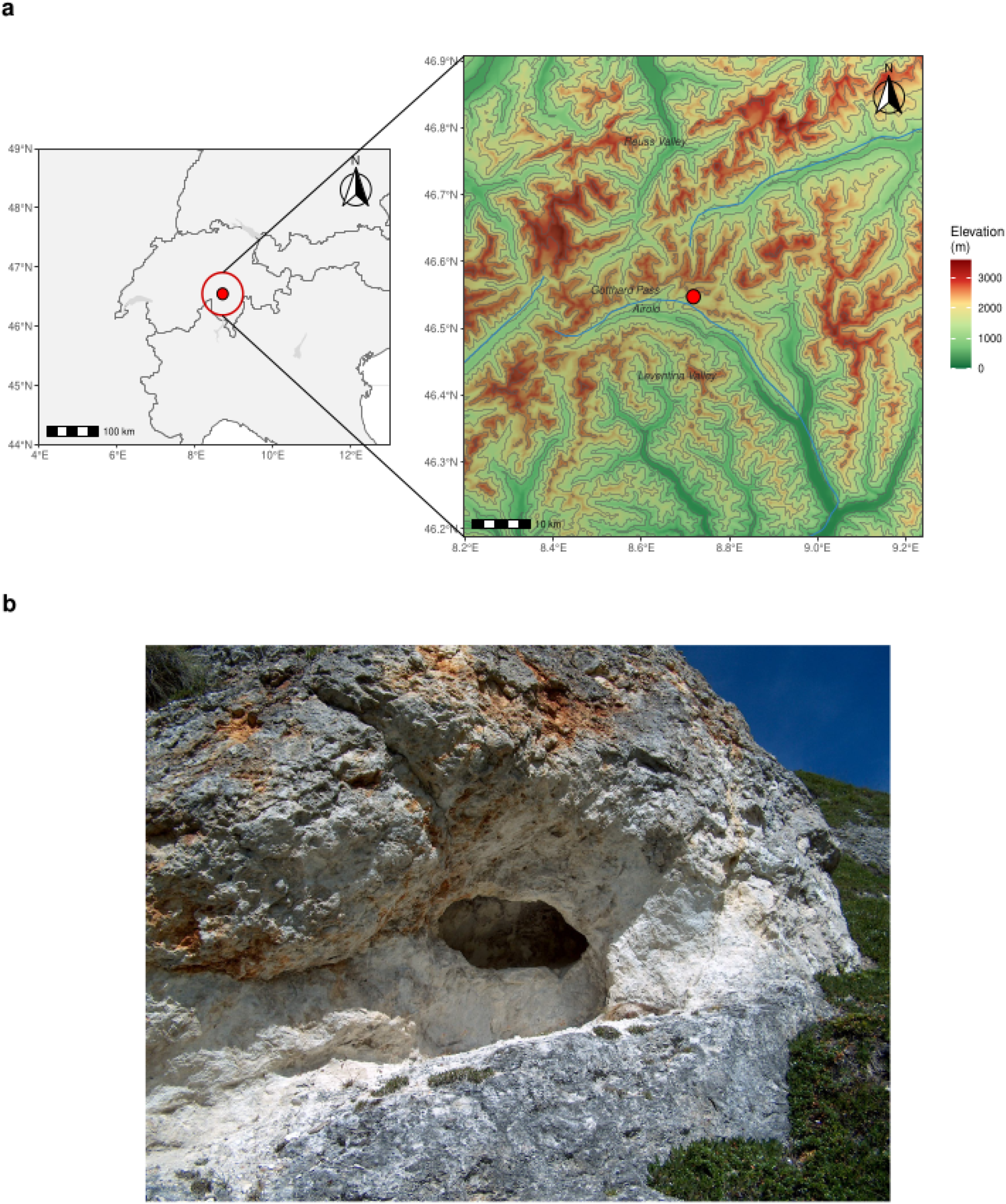
Location and topographic setting of the sampling site in the Swiss Alps. **(a)** Overview and detailed map (inset) of the sampling site, including elevation and major geographic features. A red circle indicates the sampling site. Elevation is shown as a shaded digital elevation model with 500 m contour intervals. **(b)** Field photograph of the cave entrance showing the sampling environment and surrounding habitat characteristics.

For DNA extraction, four rock samples (Rock2, Rock3, Rock4, and Rock5) containing distinct endolithic bands were selected. Before DNA isolation, the samples were scraped and crushed using a sterile metal spatula in order to separate the mineral part from the greenish organic part. Approximately 120 mg of greenish organic material from the endolithic zones was processed for DNA extraction using the DNeasy PowerSoil Pro Kit (QIAGEN, Germany), following the manufacturer’s instructions. DNA quantity and quality (A260/A280 and A260/A230) were assessed using a NanoVue spectrophotometer (Thermo Fisher Scientific, USA).

Metagenomic sequencing was performed using both Illumina short-read and Oxford Nanopore Technology (ONT) long-read platforms. Illumina libraries were sequenced on Illumina NovaSEQ6000 platform, generating 14–16 Gb of 150 bp paired-end reads for each sample. Long-read sequencing was performed using a MinION flow cell. To increase the recovery of Gloeobacterales genomic sequences from the metagenomic samples, ONT adaptive sampling [17] was performed. Five publicly available Gloeobacterales genomes were selected as target references for adaptive enrichment: *Gloeobacter violaceus, G. lilacinus, G. morelensis, G. kilaueensis*, and *Anthocerotibacter panamensis*. These genomes were used to define target regions of interest.

### Metagenome assembly

The metagenomes were first assembled from Illumina short-read sequences using the Genome-assembly pipeline implemented in the GEN-ERA toolbox v3.0.0 [18]. Briefly, raw reads were quality-filtered and trimmed using fastp v0.23.1 [19], followed by *de novo* metagenome assembly using either SPAdes v3.15.3 [20] or MEGAHIT v1.2.9 [21]. Quality-filtered reads were then mapped back to the assembled contigs using the bwa-mem algorithm in bwa v0.7.17 [22], and the resulting alignments were sorted and indexed using samtools v1.13 [23]. Metagenome binning was subsequently performed using both MetaBat2 v2.15.6 [24] and CONCOCT v1.1.0 [25]. We combined Rock2 and Rock3 during the genome assembly due to the close proximity of the two sampled rocks. Hybrid assemblies incorporating both Illumina and ONT reads were generated using the same Genome-assembly pipeline, with Flye v2.19.b1774 [26] used for long-read metagenome assembly and Pilon v1.24 [27] for polishing. Contaminant contigs were identified and removed from the hybrid assembly using Anvi’o v7.1 [28].

### Comparative metagenomic analyses

Medium-quality (MQ) metagenome-assembled genomes (MAGs), defined in this study as having CheckM completeness >50% and contamination <10%, were identified using CheckM v1.2.2 [29] and GUNC v1.0.5 [30]. Redundant genomes within each sample were dereplicated using dRep v3.0.0 [31], resulting in a final dataset of 138 MQ MAGs. These genomes were further classified using GTDB-Tk v2.0.0 [32] in the GEN-ERA toolbox [18]. Genome-wide average nucleotide identity (ANI) among the five *Gloeobacter* species was calculated using fastANI v1.33 [33], while pairwise BLASTn comparisons between the *G. violaceus* reference genome and the three other *Gloeobacter* species were visualized using BRIG v1.1.1 [34]. Pairwise ANI and average amino-acid identity (AAI) values of the 15 cyanobacterial MAGs were also calculated using fastANI and FastAAI v0.1.20 [35] against 9,026 cyanobacterial/melainabacterial genomes from NCBI Genome Datasets [36]. The five *Gloeobacter* genomes were annotated with Prokka v1.15.6 [37], and orthologous gene groups were inferred using OrthoFinder v2.5.4 [38]. Functional annotation of predicted protein sequences into Clusters of Orthologous Groups (COG) categories was performed using COGclassifier v2.0.0 (https://github.com/moshi4/COGclassifier). We retrieved hidden Markov model profiles (pHMMs) for genes of interest (e.g., associated with phycobilisomes, photosynthesis, and respiratory machinery) from the Kyoto Encyclopedia for Genes and Genomes (KEGG) database [39] and used hmmsearch in HMMER v3.4 (http://hmmer.org/) to identify their homologs in Gloeobacterales genomes.

### Phylogenomic and divergence time analyses

We combined 15 high-quality (HQ) cyanobacterial MAGs with CheckM completeness >80% with 254 representative Cyanobacteriota genomes (Table S1) selected from the curated dataset of Sudianto et al. [40] and Goremykin et al. [41]. We used GToTree v1.8.8 [42] to generate a phylogenomic supermatrix alignment from 251 individual alignments that were aligned and trimmed using MUSCLE v5.1 [43] and TrimAl v1.4.rev15 [44], respectively. The phylogenomic tree was reconstructed using IQ-TREE v3.1.0 [45] with automated model selection using ModelFinder (-m MFP; [46]) and 1,000 ultrafast bootstrap replicates [47].

Divergence times were estimated using three calibration points from the cyanobacterial phylogeny of Boden et al. [48], based on geochemical and fossil evidence: (1) the diversification of cyanobacteria, constrained between 2.32 billion years ago (Ga) to 3 Ga [49,50]; (2) the origin of cyanobacteria possessing apical cells, constrained between 1.7 to 1.888 Ga [51], and (3) the origin of cyanobacteria with akinetes, constrained between 1.6 to 1.888 Ga [52]. The root age was additionally constrained to <4.1 Ga, following the proposed maximum age of stem cyanobacteria in Sánchez-Baracaldo et al. [53]. Divergence time estimation was performed using the IQ2MC workflow in IQ-TREE [54] together with the approximate likelihood method implemented in MCMCTree [55] from PAML v4.10.8.iq2mc [56]. Briefly, an alignment of 41 small and large ribosomal proteins from the 265 taxa (four were removed due to identical sequences), together with the constrained topology obtained from the phylogenomic analysis, was used to estimate the branch-length gradients and Hessians required by MCMCTree. These estimates were subsequently used to infer divergence times under the following parameters: “usedata = 2, clock = 3, sampfreq = 100, nsample = 40000, and burnin = 1000000”. Two independent Markov chain Monte Carlo (MCMC) runs were performed, and convergence was assessed using Tracer v1.7 [57] by verifying that all effective sample sizes (ESS) exceeded 200. The tree was visualized using iTOL v7 [58].

### Identification of cyanase and cyanate transporter genes

To identify cyanase (*cynS*) and cyanate substrate-binding protein (SBP) homologs, we used pHMMs corresponding to *cynS* (KEGG Orthology K01725) and the NMT1-like family (Pfam PF13379), respectively. The *cynS* pHMM was searched using hmmsearch against GTDB release R220 (113,104 bacterial and archaeal representative genomes; [59]), all publicly available Gloeobacterales genomes, and all HQ cyanobacterial MAGs in this study. Hits with an *E*-value ≤ 1 × 10^-5^ were retained. The NMT1-like pHMM was searched against all cyanobacterial MAGs reconstructed in this study, and the resulting SBP dataset was supplemented with BLASTp hits from *Gloeobacter dolomitensis* and the characterized *Synechococcus elongatus* PCC 7942 proteins NrtA (YP_400256), CmpA (YP_400505), and CynA (AAB58736). The identified protein sequences were aligned using MAFFT v7.5.2 [60] and trimmed with ClipKIT v2.4.1 using default parameters [61].

Maximum-likelihood gene trees were reconstructed using IQ-TREE v3.1.0 [45] with the LG+F+G substitution model and 1,000 ultrafast bootstrap replicates. Trees were formatted and annotated using the format-tree.pl function in Bio-MUST-Core v0.252040 (https://metacpan.org/dist/Bio-MUST-Core), and visualized using iTOL [58].

Structural similarity of the *G. dolomitensis* potential cyanate SBP to the identified SBP homologs was assessed using US-align v20260527 in default parameters [62]. The *G. dolomitensis* potential cyanate SBP structure was first predicted using AlphaFold Server [63] and then structurally aligned against reference proteins from the RCSB Protein Data Bank [64], including nitrate-, bicarbonate-, taurine-, and alkanesulfonate-binding proteins. Because no experimentally determined structures of cyanate-binding SBPs are currently available, we additionally predicted the structures of CynA from *Anthocerotibacter panamensis* and *Synechococcus elongatus* PCC 7942 using AlphaFold Server. These predicted structures were then included in the structural comparisons with the *G. dolomitensis* SBP.

### Horizontal gene transfer inference

Horizontal gene transfer (HGT) events among phyla, classes, orders, families, and genera within each microbial community were inferred using MetaCHIP v1.10.10 [65] with the “-r pcofg” option, using taxonomic assignments obtained from GTDB-Tk. To visualize HGT events involving cyanobacteria, only transfers in which either the donor or recipient belonged to cyanobacteria were retained. These interactions were visualized as chord diagrams using circlize v0.4.18 [66] in R.

## Results

### Taxonomic composition differs among neighboring endolithic communities

We compiled all MQ MAGs and classified them using the GTDB taxonomy. Despite the close spatial proximity of these rock communities, the recovered bacterial communities differed substantially in taxonomic composition (Figure 2a). Actinomycetota represented the dominant phylum in Rock2-3, whereas Pseudomonadota predominated in both Rock4 and Rock5. The second most abundant phyla also differed between samples, with Planctomycetota and Cyanobacteriota enriched in Rock4 and Actinomycetota in Rock5. In total, we assembled 138 MQ MAGs across the three datasets, 20 of which were classified as cyanobacteria.

**Figure 2.**
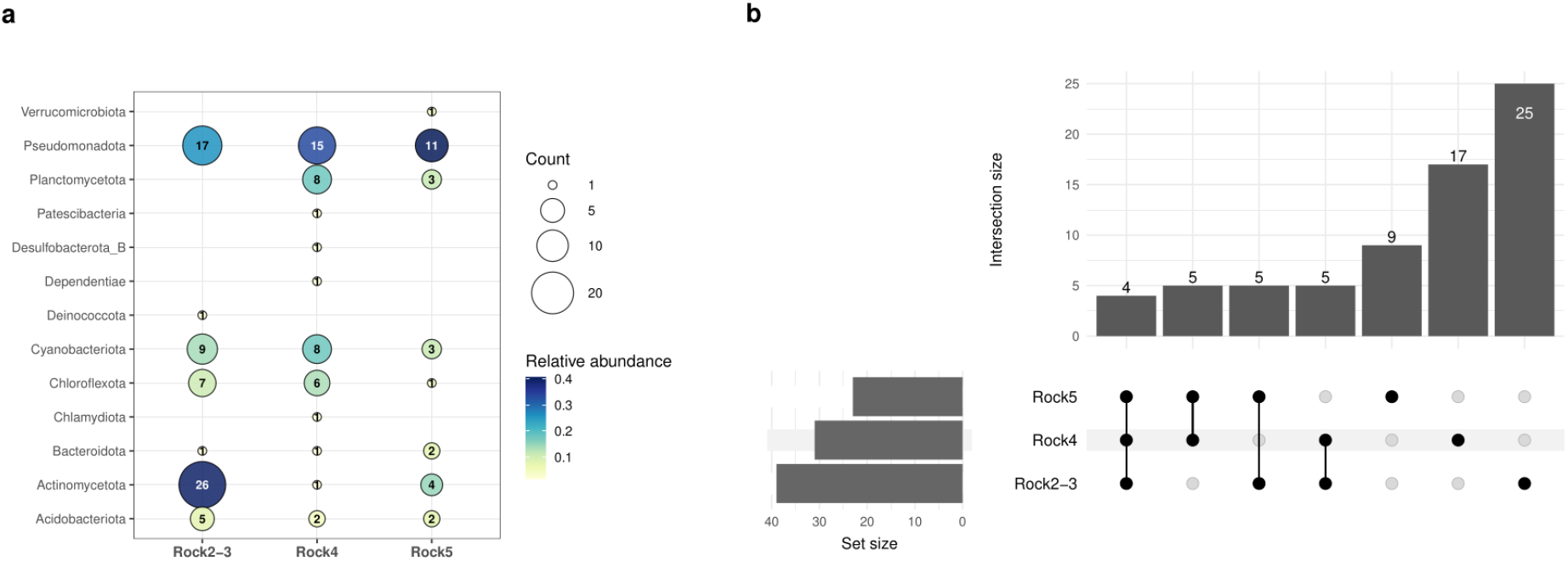
Diversity of medium-quality (MQ) metagenome-assembled genomes (MAGs) recovered from the three rock samples. **(a)** Number of MQ MAGs assigned to each phylum across the three rock samples. The colored sectors within each circle indicate the relative abundance of each phylum in the corresponding rock sample. **(b)** UpSet plot showing the number of shared and unique MQ MAGs among the three rock samples at the family level.

Among the 138 MQ MAGs, only four families were shared across all three communities (Figure 2b): Gloeobacteraceae and Elainellaceae (Cyanobacteriota), Pseudonocardiaceae (Actinomycetota), and Sphingomonadaceae (Pseudomonadota). Taxonomic overlap was even more limited at the species level, where only a single *Gloeobacter* genome was recovered from all three samples. This genome could not be assigned to any described species by GTDB, indicating that it may represent a previously uncharacterized *Gloeobacter* species.

### The Piora Valley dolomite rocks host novel and diverse cyanobacteria

We reconstructed a phylogenomic tree comprising the 15 HQ cyanobacterial MAGs recovered from the dolomite rock samples, along with 254 representative Cyanobacteriota genomes (Figure 3; Figure S1). The 15 MAGs were distributed across six order-level lineages: Gloeobacterales, FACHB-261, Acaryochloridales, Oculatellales, Chroococcidiopsidales, and Nostocales. Pairwise ANI comparisons between these 15 cyanobacteria and 9,026 cyanobacterial/melainabacterial genomes indicated that nearly all represent potentially novel species, with 14 of 15 MAGs exhibiting maximum ANI values ranging from 79% to 94% (Table S2). Pairwise AAI comparisons further suggested that three MAGs may represent lineages with potential genus-level novelty, including Rock2-3_bin-18 (Acaryochloridales) (GCA_041743695.1; AAI: 64.73%), Rock2-3_bin-47 (Oculatellales) (GCA_031374485.1; AAI: 59.19%), and Rock4_megahit-12 (Chroococcidiopsidales) (GCA_023558375.1; AAI: 63.6%).

**Figure 3.**
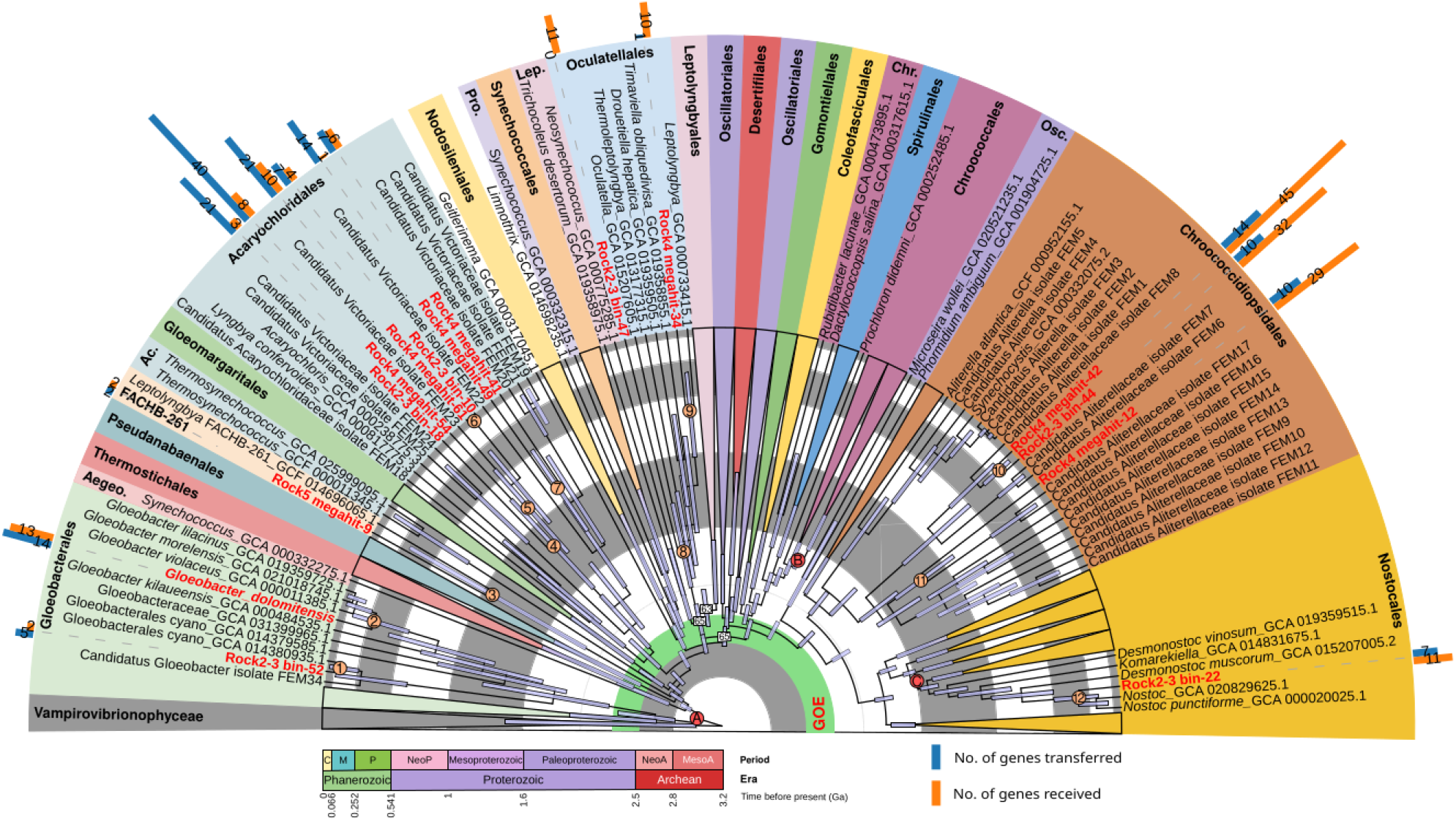
Time-calibrated phylogeny and horizontal gene transfer history across cyanobacterial genomes. A dated phylogenetic tree inferred from the phylogenetic tree in Figure S1 and three calibration points (A–C). The horizontal bars at the nodes represent the 95% highest posterior density (HPD) intervals. All bootstrap supports are >90%, except for labelled nodes. Nodes of interest are labeled from 1–12. The numbers below the period and era legend represent the time in billion years ago (Ga). New genomes reported in this study are marked in red. The accompanying bar graph summarizes the number of horizontal gene transfer (HGT) events (transfered/received) predicted for each of the new genomes in this study. GOE, Great Oxygenation Event; Aegeo., Aegeococcocales; Ac., Acaryochloridales; Pro., Prochlorotrichales; Lep., Leptolyngbyales; Chr., Chroococcales; Osc., Oscillatoriales; C., Cenozoic; M., Mesozoic; P., Paleozoic; NeoP., Neoproterozoic; NeoA., Neoarchean; MesoA., Mesoarchean.

Only Rock5_megahit-9 showed >95% ANI to the MAG “*Leptolyngbya*” sp. SOY_34_metabat_272 (GCA_031399305.1), recovered from a glacier stream in Valsorey, Switzerland. In the phylogenomic tree, Rock5_megahit-9 was placed as a sister to “*Leptolyngbya*” FACHB-261 (Figure 3). Together, these genomes formed a distinct clade positioned between Pseudanabaenales and Gloeomargaritales—consistent with the distinct order-level lineage currently designated as FACHB-261 in GTDB. The inclusion of Rock5_megahit-9 therefore expands the genomic representation of this lineage and provides additional evidence for its distinctiveness. However, formal taxonomic validation and the establishment of a validly published name for this genus and order would require additional evidence, including broader taxon sampling and assessment of its phylogenetic stability.

The most abundant cyanobacterial MAGs we identified here were broadly classified as “*Ca*. Victoriaceae” (six MAGs) and “*Ca*. Aliterellaceae” (four MAGs). These two clades are family-level taxa recently reported in Goremykin et al. [41], recovered from Antarctic endolithic rocks. Overall, the Piora Valley dolomite rocks harbored a phylogenetically diverse cyanobacterial community dominated by lineages that remain poorly represented in current genomic databases.

### A novel *Gloeobacter* species from dolomite rocks harbors a cyanate-associated genomic locus

Among the diverse cyanobacterial lineages recovered from the Piora Valley dolomite, the *Gloeobacter* genome shared across all rock communities was particularly notable because it could not be assigned to any described species. To improve the contiguity of this putative novel *Gloeobacter* genome, we generated a hybrid assembly incorporating long-read sequencing data. After contaminant removal, the resulting MAG comprised 87 contigs with a total genome size of 4.8 Mbp and estimated CheckM completeness and contamination of 97.44% and 5.13%, respectively (Table S3). Its genome size and GC content were within the ranges reported for the other four available *Gloeobacter* genomes (4.6–4.9 Mbp and 60–62%, respectively; Table S3). Pairwise ANI values between the novel MAG and the four available *Gloeobacter* genomes ranged from 78 to 89%, well below the commonly accepted species threshold, supporting its classification as a putative novel species (Figure 4a). Following the SeqCode [67] and recommended guidelines for description of new cyanobacterial species [68], we propose the name *Gloeobacter dolomitensis* for this lineage.

**Figure 4.**
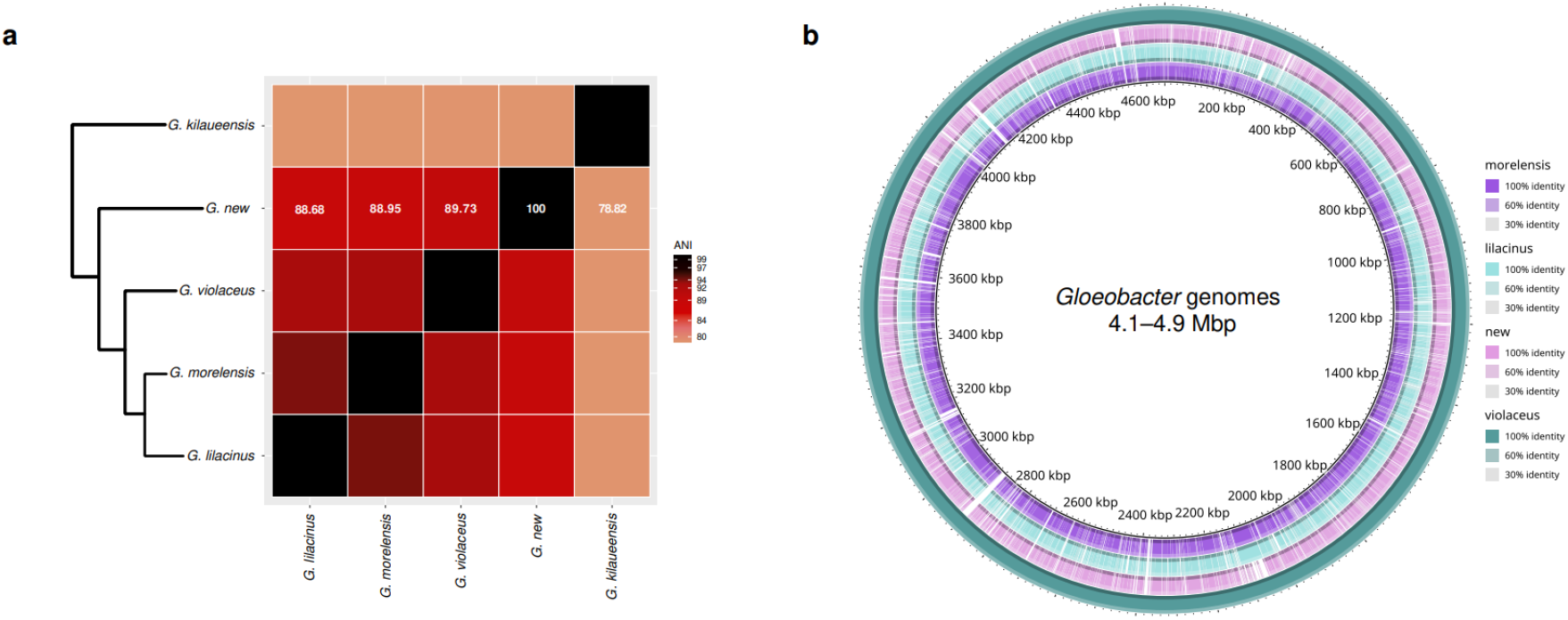
Comparative genomic analysis of the novel *Gloeobacter* species and previously published *Gloeobacter* genomes. **(a)** Genome-wide average nucleotide identity (ANI) among the five *Gloeobacter* genomes. ANI values between *G. dolomitensis* and the other *Gloeobacter* genomes are shown in white. The phylogenetic backbone is adapted from Figure S1. **(b)** BLASTn-based whole-genome comparison of *G. morelensis, G. lilacinus*, and *G. dolomitensis* against the *G. violaceus* genome, visualized using BRIG.

Pairwise BLASTn analysis revealed extensive regions of high sequence similarity between *G. dolomitensis* and *G. violaceus*, interspersed with localized regions of sequence divergence (Figure 4b). The *G. dolomitensis* genome possesses a gene repertoire for phycobilisomes, photosynthesis, and respiratory machinery that is largely consistent with those of other *Gloeobacter* genomes, except for bacteriorhodopsin, which is unique to *G. violaceus* among the genus (Figure S2). Orthogroup analysis further showed that approximately 56% of the orthogroups (OGs) are conserved across the five analyzed *Gloeobacter* genomes (Figure S3). In contrast, only 53 orthogroups (81 genes) were unique to the novel *Gloeobacter* genome. Functional annotation assigned COG categories to 64 of these genes, with the largest representation in categories related to the mobilome, transcription, and cell wall/membrane/envelope biogenesis (Figure S4; Table S4).

Among these lineage-specific genes, one showed a strong homologous match to *cynS* (*E*-value = 2.74 × 10^-99^; 60.7% identity), encoding cyanase, which catalyzes the degradation of cyanate. The gene has an unusual distribution within Gloeobacterales, occurring in *G. dolomitensis* and several other Gloeobacterales genomes (10 of 18) but being absent from the other examined *Gloeobacter* species (Figure 5a). Notably, in four Gloeobacterales genomes (including *Anthocerotibacter*), *cynS* is adjacent to a potential ABC transporter operon (complete or partial) comprising a cyanate SBP (*cynA*), a permease (*cynB*), and an ATP-binding protein (*cynD*; Figures 5a and 5b). The SBP close to *cynS* in *Anthocerotibacter* and two other Gloeobacteraceae cluster phylogenetically with characterized CynA proteins, supporting their assignment as cyanate-binding components of the CynABD transporter (Figure 5c).

**Figure 5.**
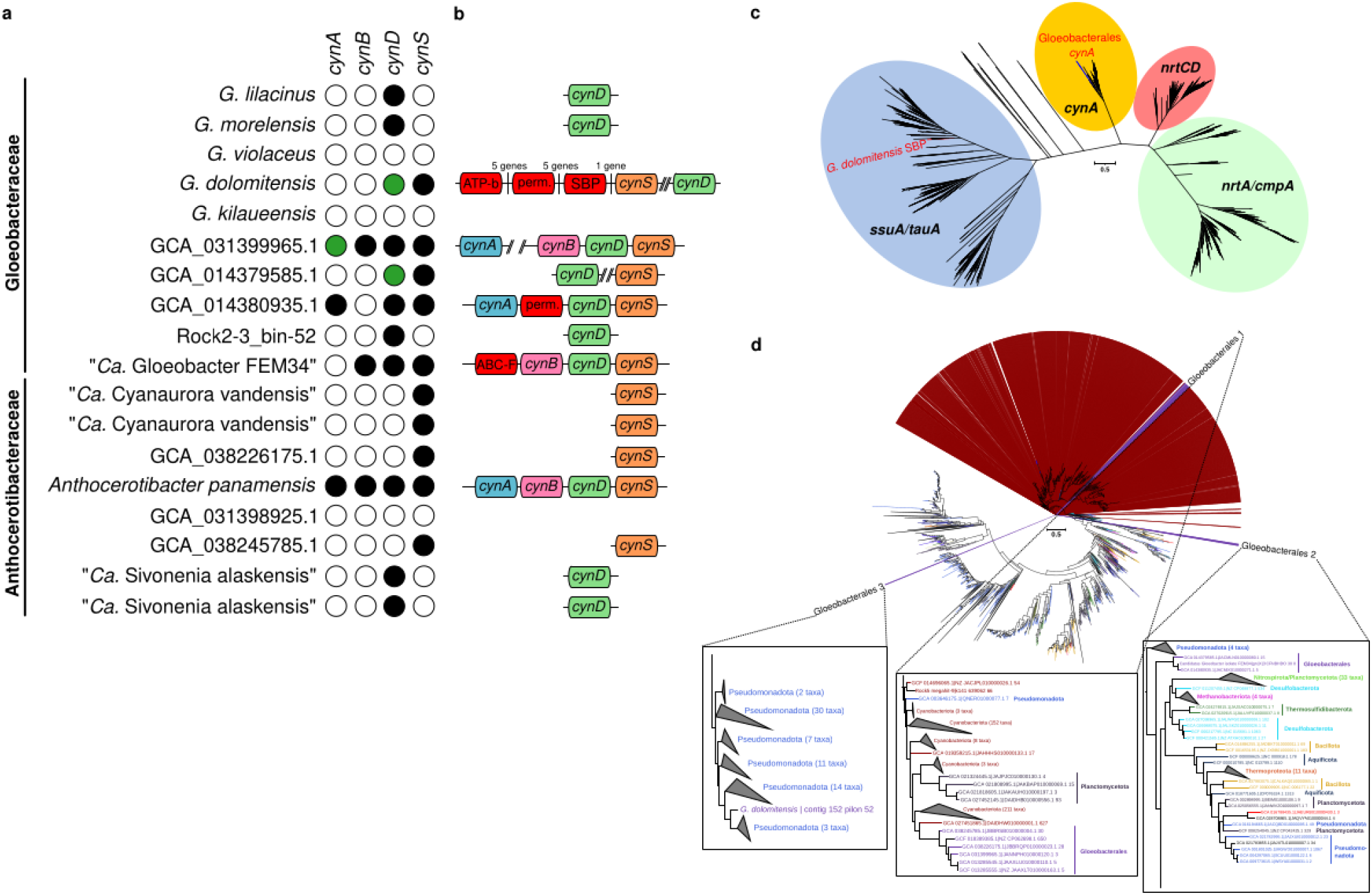
Distribution and evolution of cyanase and cyanate transporter genes among the Gloeobacterales. **(a)** Presence of genes encoding cyanate transporter SBP (*cynA*), permease (*cynB*), and ATP-binding protein (*cynD*) across the Gloeobacterales. **(b)** Organization of the *cynABDS* locus in Gloeobacterales genomes. Red boxes indicate non-orthologous genes occupying the typical *cynABD* locus, and numbers in *G. dolomitensis* indicate the number of intervening genes. **(c)** Gene tree of nitrate/bicarbonate/cyanate/sulfonate/taurine-binding proteins. Gloeobacterales *cynA* sequences and the *G. dolomitensis* SBP are highlighted in red. Ellipses in the gene tree denote sequences with diverse substrate specificity. **(d)** Gene tree of cyanase (*cynS*). Gloeobacterales and other cyanobacterial *cynS* sequences are highlighted in purple and maroon, respectively. Insets show the three Gloeobacterales *cynS* clusters; font colors indicate bacterial groups independently of the ellipse color scheme in panel (c). Maximum-likelihood trees were reconstructed using LF+F+G substitution models and 1,000 ultrafast bootstrap replicates in the following alignment dimensions: CynA (1,338 sequences × 1,301 sites) and CynS (8,358 sequences × 1,302 sites). SBP, substrate-binding protein; perm., permease; ATP-b, ATP-binding protein; ABC-F, ABC-F family ATP-binding cassette domain-containing protein; *ssuA*, aliphatic sulfonate-binding protein; *tauA*, taurine-binding protein; *nrtA*, nitrate/nitrite-binding protein; *cmpA*, bicarbonate-binding protein; *nrtCD*, ATP-binding protein of nitrate/nitrite ABC transporter.

In *G. dolomitensis*, however, *cynS* is associated with a non-canonical ABC transporter comprising a sulfonate/taurine-binding protein-like SBP, an Acidobacteria-like permease, and a LolD-like ATP-binding protein (Figure 5b). Structural prediction (Table S5) and gene tree (Figure 5c) placed the SBP with highest similarity to aliphatic sulfonate- or taurine-binding proteins (template modeling score (TM)-score = 0.87–0.88; root-mean-square deviation (RMSD) = 2.1–2.5 Å). Although these similarities suggest that the protein belongs to a broader class of ligand-binding proteins that includes sulfonate/taurine-binding SBP, they do not establish its substrate specificity. The close genomic association between this transporter and *cynS*, together with the lack of a canonical CynABD transporter, raises the possibility that this divergent ABC transporter contributes to cyanate uptake. However, direct evidence for cyanate transport is currently lacking.

Phylogenetic analysis (Figure 5d) indicated that *cynS* of Gloeobacterales can be divided into three groups: (1) clustered within cyanobacteria (all Anthocerotibacteraceae and one Gloeobacteraceae); (2) sister to diverse archaea and bacteria (three Gloeobacteraceae); and (3) clustered within Pseudomonadota (*G. dolomitensis*). This result is consistent with multiple horizontal acquisitions of *cynS* within Gloeobacterales, namely from other cyanobacteria, Pseudomonadota, or unknown sources. Together, the presence of horizontally acquired *cynS* and its association with distinct ABC transporter architectures compared to the other Gloeobacterales support the hypothesis that cyanate-related metabolisms may occur in *G. dolomitensis* and raise the possibility that cyanate uptake in this lineage involves a non-canonical transporter system.

### Gloeobacterales shows a strong association with interphylum horizontal gene transfer

To characterize gene exchange within the endolithic microbiomes, we inferred HGT among all MQ MAGs recovered from the three rock communities. In total, 1,163 HGT events were inferred among 138 genomes (Table S6). HGT was strongly biased toward transfers between closely related organisms, with ∼74% (862/1163) of all inferred events occurring within the same bacterial phylum (Table S6; Figure S5–S7).

Cyanobacterial lineages displayed marked differences in their HGT profiles (Figure 3; Figure S8). Acaryochloridales represented the dominant source of horizontally transferred genes, accounting for ∼63% (110/173) of all cyanobacterial donor events (Figure S8; Table S7). Conversely, Chroococcidiopsidales primarily acted as recipients, receiving ∼57% (106/187) of cyanobacterial HGT events while contributing only ∼19% (34/173) of donor events (Figure S8; Table S7). Interphylum HGT represented only ∼10% (39/360) of all inferred transfers in cyanobacteria (Figure S8). Nevertheless, Gloeobacterales exhibited a disproportionately high frequency of interphylum interactions compared with other cyanobacterial orders, with ∼72% (24/33) of HGT events involving inferred donor or recipient relationships with non-cyanobacterial taxa (Table S8).

Together, these results reveal pronounced asymmetry in HGT contributions among cyanobacterial lineages: Acaryochloridales primarily acted as donors, whereas Chroococcidiopsidales were predominantly recipients. Gloeobacterales, in contrast, showed a comparatively strong association with interphylum gene exchange. These lineage-specific HGT profiles indicate that gene exchange was unevenly distributed across the cyanobacterial community, with different lineages exhibiting distinct inferred donor, recipient, and interphylum-interaction profiles.

### Endolithic cyanobacteria comprise phylogenetically diverse lineages with divergence times spanning the Proterozoic and Phanerozoic

The 15 endolithic cyanobacterial MAGs exhibited substantial variation in inferred divergence times, spanning approximately 20 million years ago (Ma) to 1.68 Ga (Figure 3; Table S9). Rather than being restricted to a single temporal interval, the MAGs were distributed across both Phanerozoic and Proterozoic eras. The oldest lineage, Rock2-3_bin-47, was estimated to have diverged 1.68 Ga (95% Highest Posterior Density (HPD): 1.28–1.98 Ga; Node 8). This lineage is also phylogenetically distinct, forming a sister lineage to the remaining sampled Oculatellales, consistent with its potential status as a novel genus (Table S2). Two additional lineages with potential novel genus-level status had mean divergence estimates of approximately 1.23 Ga and 1.22 Ga (95% HPDs: 0.72–1.68 and 0.89–1.47 Ga, respectively; Nodes 4 and 11), placing their inferred divergences within the Mesoproterozoic. A fourth lineage had a mean estimate of 1.08 Ga, although its broad HPD (0.32–1.94 Ga; Node 3) indicates substantially greater temporal uncertainty, possibly reflecting limited taxon sampling within the FACHB-261 lineage.

Several additional endolithic lineages had mean divergence estimates during the Neoproterozoic, ranging from approximately 0.66 to 0.88 Ga and including representatives of Acaryochloridales and Oculatellales. The remaining MAGs were associated with the Phanerozoic era, with mean estimates ranging from 20 to 328 Ma. Within this younger group, the *Gloeobacter* lineage represented by *G. dolomitensis* was estimated to have diverged at 289.9 Ma (95% HPD: 115.0–581.4 Ma; Node 2). Overall, eight of the 15 MAGs had mean divergence estimates older than 500 Ma, including four with mean estimates exceeding 1 Ga, demonstrating that the endolithic cyanobacterial lineages recovered from the Piora Valley encompass a broad range of evolutionary ages.

The temporal distribution of these MAGs mirrors their phylogenetic dispersion across the cyanobacterial tree, with endolithic representatives occurring at multiple evolutionary depths rather than forming a single temporally coherent lineage. Thus, the dated phylogeny identifies multiple endolithic lineages with Proterozoic divergence estimates, while also revealing substantially younger endolithic lineages that originated during the Phanerozoic.

### Description of *Gloeobacter dolomitensis* sp. nov

*Gloeobacter dolomitensis* (do.lo.mi.ten′sis. (N.L. masc. adj.), N. L. n. *dolomites*, dolomite rock; N. L. masc. adj. ending *-ensis*, indicating association with a substrate or habitat; N. L. masc. adj. *dolomitensis*, inhabiting or associated with dolomite rock). We reconstructed the type genome of *G. dolomitensis* from metagenomic sequencing data from a dolomite-associated endolithic microbial community. The genome is 4.8 Mb, with estimated CheckM completeness and contamination of 97.44% and 5.13%, respectively. We recovered the genome from sequencing reads generated from endolithic dolomite rocks collected from the Piora Valley in the southern Swiss Alps. Phylogenomic analyses place *G. dolomitensis* within the genus *Gloeobacter* and the phylum Cyanobacteriota. The closest available genome is *G. violaceus*, showing an ANI of 89.73%, supporting its recognition as a distinct species. The genome encodes 98 of the 126 genes associated with oxygenic photosynthesis, phycobilisome assembly, and respiratory metabolism. The type genome of *G. dolomitensis* is deposited in the NCBI under genome accession number JCENRC000000000. *G. dolomitensis* belongs to the family Gloeobacteraceae, order Gloeobacterales, class Cyanophyceae, and phylum Cyanobacteriota. The name has been registered under SeqCode: https://seqco.de/i:56654.

## Discussion

Previous studies of the Piora Valley dolomite rocks identified cyanobacteria as important components of the endolithic phototrophic community, including representatives related to *Leptolyngbya, Nostoc, Scytonema, Anabaena, Spirirestis, Microcoleus, Chroococcidiopsis*, and *Gloeobacter* [7,8]. Our shotgun metagenomic approach substantially expands this view, recovering 15 high-quality cyanobacterial MAGs representing previously unrecognized genus- or species-level diversity (Figure 3; Table S2). Rock5_megahit-9 is closely related to a previously published but formally undescribed genome [69] and may represent an early-diverging lineage within the recently proposed FACHB-261 order [70]. Notably, only a single *Gloeobacter* species was recovered across all sampled rocks, which we describe here as the novel species *G. dolomitensis*.

The substantial taxonomic novelty observed among the 15 cyanobacterial MAGs highlights the extent to which endolithic communities remain underrepresented in genomic surveys. Three MAGs showed divergence consistent with potential genus-level novelty, while most of the remaining MAGs showed species-level divergence from available reference genomes. Several Piora Valley MAGs also clustered with “*Ca*. Victoriaceae” and “*Ca*. Aliterellaceae” (Figure 3), previously recovered from Antarctic endolithic rocks [41]. Their occurrence across geographically distant endolithic habitats is consistent with broad dispersal or repeated colonization, although broader sampling is needed to establish habitat associations. At the same time, the limited taxonomic overlap between Piora Valley and endolithic communities from the deserts of Atacama, Negev, Mojave, Al-Jafr Basin, and other environments [71–75] suggests substantial habitat-specific turnover. Elevation and geology likewise influence community composition in Antarctic endolithic systems [76], suggesting that the distinctive Piora Valley assemblage may reflect the combined influence of alpine climate and dolomite geology.

The recovery of multiple Gloeobacterales genomes from Piora Valley is particularly notable given the sparse genomic representation of this lineage. We discovered two previously uncharacterized Gloeobacterales genomes, including *G. dolomitensis* and the more deeply branching Rock2-3_bin-52. This is notable given the sparse genomic representation of Gloeobacterales, which currently comprises fewer than 20 genomes in NCBI and GTDB (last accessed 20 August 2026). Gloeobacterales is the only sister lineage of remaining extant photosynthetic cyanobacteria and characterized by the absence of thylakoid membranes [9,10]. Its apparent rarity in genomic databases may partly reflect low abundance, slow growth, and historical sampling bias [13]. The recovery of multiple Gloeobacteraceae lineages from a single endolithic habitat suggests that the diversity of this poorly sampled lineage is substantially underestimated.

The recurring presence of *G. dolomitensis* across various samples in Piora Valley also provides an opportunity to examine metabolic features that may contribute to its persistence in a nutrient-limited endolithic environment. The endolithic environment imposes desiccation, freeze–thaw cycles, nutrient limitation, variable irradiance, and UV-B exposure [3,77], conditions that can strongly constrain microbial growth and favor stress tolerance and metabolic flexibility. These stresses resemble those encountered in other extreme environments, including the Arctic and Antarctic [71]. Nitrogen availability may be an additional constraint in this environment, with total nitrogen measured at only 0.02% (w/w) in the Piora Valley dolomite rocks [7].

Here we show that the *G. dolomitensis* genome contains a *cynS* gene that encodes cyanase (Figure 5; Table S4). *CynS* catalyzes the decomposition of cyanate in the presence of bicarbonate to CO_2_ and ammonium, thereby providing a potential route for recovering nitrogen from cyanate [78]. Cyanate utilization has been experimentally demonstrated in cyanobacteria, including *Synechococcus elongatus* PCC 7942, in which the *cynABDS* locus encodes a dedicated cyanate-utilization system [78,79]. In *Synechococcus*, CynA functions as the periplasmic substrate-binding component of an ABC transporter, and disruption of *cynA* abolishes active cyanate transport, demonstrating that uptake is an integral component of cyanate utilization [79]. The presence of *cynS* therefore suggests that *G. dolomitensis* has the potential to degrade cyanate, potentially as an alternative nitrogen source and/or as a detoxification mechanism [80].

The genomic context of *cynS* provides additional evidence for an unusual cyanate-related system in *G. dolomitensis*. In *Anthocerotibacter* and two other Gloeobacteraceae genomes, *cynS* is associated with a canonical CynA-like SBP and the corresponding permease and ATP-binding components, consistent with the organization of characterized cyanate transport systems (Figures 5a–c). This finding echoes a previous study [79] that highlights the substrate-binding and permease proteins located near *cynS* as a cyanate transporter are frequently misannotated as nitrate transporter proteins.

In *G. dolomitensis*, however, *cynS* is located near to a distinct ABC transporter whose SBP contains the PF13379/NMT1-like domain but is phylogenetically distinct from CynA (Figure 5c). This observation reflects the broader structural relationship among NrtA/CmpA-, SsuA-, and TauA-like SBP, which share a conserved substrate-binding architecture despite differences in ligand specificity [81,82]. The functional diversity within this broader transporter family is further illustrated by CynABD of *Synechococcus elongatus*, which can transport both cyanate and nitrite [83]. This indicates that substrate assignments based solely on sequence similarity or structural classification may not fully capture the substrate range of related ABC transport systems. Nevertheless, the structural and phylogenetic similarity of the *G. dolomitensis* SBP to other ligand-binding proteins does not establish cyanate specificity, and direct functional characterization will be required.

The association of this divergent ABC transporter with *cynS* remains noteworthy. We hypothesized that the canonical *cynABDS* operon was lost in the ancestor of the *Gloeobacter*, as they are present in several Gloeobacterales taxa but not *Gloeobacter* (Figure 5a). *G. dolomitensis* then likely recruited or retained a non-orthologous transporter capable of recognizing cyanate, potentially through broader substrate specificity. Alternatively, the transporter may recognize a different substrate, with its proximity to *cynS* reflecting genomic rearrangement or co-localization of genes involved in nitrogen or sulfur metabolism rather than a direct functional relationship.

Indeed, only 12 out of 55 marine picocyanobacterial genomes encode *cynS* with the *cynABD* transporter [84]. The frequent occurrence of *cynS* without an associated *cynABD* locus has led to the hypothesis that cyanase may also function primarily in cyanate detoxification, including the degradation of cyanate generated as a byproduct of cellular metabolism, such as urea-cycle or carbamoyl-phosphate-related reactions [78,84]. We observed the same pattern in Gloeobacterales, with only 4 of the 10 *cynS*-containing genomes having both *cynS* and either a complete or partial *cynABD* operon (Figure 5a). Thus, the *G. dolomitensis* locus represents an unusual genomic configuration in which a horizontally acquired *cynS* is associated with a distinct, non-CynA-like ABC transporter. The available genomic and structural evidence does not demonstrate that this transporter imports cyanate, but its co-localization with *cynS* provides a testable hypothesis for a either a potentially non-canonical route of cyanate uptake or a genomic association unrelated to cyanate transport.

The evolutionary history of *cynS* further supports a scenario of horizontal acquisition of cyanate-degradation capacity. Phylogenetic analysis indicates that the *G. dolomitensis cynS* clustered within Pseudomonadota, consistent with horizontal acquisition from a Pseudomonadota lineage and is distinct from the cyanobacterial *cynS* sequences associated with canonical *cynABDS* loci (Figure 5d). Together with the absence of a canonical *cynABD* transporter, these findings suggest that cyanate metabolism in *G. dolomitensis* may have evolved through a combination of horizontal acquisition of *cynS* and the recruitment, retention, or genomic association of an alternative transporter.

The recruitment of a foreign and chimeric ABC transporter has previously been reported for ribose transport in Gloeobacteraceae [40], and the propensity for HGT between *Gloeobacter* and distant bacterial phyla observed in this study (Figure 3) provides further precedent for such genomic innovation. Such metabolic flexibility could be advantageous in the nitrogen-poor Piora Valley dolomite environment, where access to alternative nitrogen sources may provide a selective benefit. Overall, the presence of *cynS* in combination with distinct transporter architectures across Gloeobacterales suggests that cyanate metabolism and its genomic organization may be more evolutionarily flexible in this lineage than previously recognized. However, the available evidence does not distinguish whether the *G. dolomitensis* system functions primarily in cyanate assimilation, cyanate detoxification, or another process. Whether *G. dolomitensis* can use cyanate as a nitrogen source, and whether the adjacent ABC transporter contributes to cyanate uptake, will require experimental validation.

## Conclusions

Our study reveals previously unrecognized cyanobacterial diversity in the Piora Valley dolomite and expands the genomic representation of the early-diverging Gloeobacterales. The recovery of *G. dolomitensis* and other divergent lineages highlights the value of genome-resolved approaches for characterizing microbial diversity in poorly sampled endolithic environments. Beyond taxonomic diversity, the genomic features identified here suggest that survival in this nutrient-limited habitat may involve substantial metabolic flexibility and evolutionary innovation. In particular, the divergent cyanate-associated system in *G. dolomitensis* illustrates how horizontal gene transfer and alternative transporter architectures may contribute to the emergence of metabolic capabilities in isolated microbial lineages. Together, these findings emphasize that endolithic habitats remain an important reservoir of underexplored cyanobacterial diversity and provide opportunities to uncover novel links between environmental constraints, metabolic adaptation, and microbial evolution.

## List of abbreviations

AAI: Average Amino-acid Identity
ABC: ATP-synthase Binding Cassette
ANI: Average Nucleotide Identity
BLASTn: Nucleotide Basic Local Alignment Search Tool
BLASTp: Protein Basic Local Alignment Search Tool
COG: Clusters of Orthologous Groups
ESS: Effective Sampling Sizes
Ga: Billion years ago
GTDB: Genome Taxonomy Database
HGT: Horizontal Gene Transfer
HMM: Hidden Markov model
HPD: Highest Posterior Density
HQ: High Quality
KEGG: Kyoto Encyclopedia for Genes and Genomes
Ma: Million years ago
MAG: Metagenome-Assembled Genomes
MCMC: Markov chain Monte Carlo
MQ: Medium Quality
NCBI: National Center for Biotechnology Information
OG: Orthologous Groups
ONT: Oxford Nanopore Technology
pHMM: Hidden Markov model profile
RMSD: Root-mean-square deviation
SBP: Substrate-binding protein
TM-score: Template Modeling score

## Declarations

### Ethics approval and consent to participate

Not applicable

### Consent for publication

Not applicable

### Availability of data and material

All supplementary data and custom scripts used in this study are deposited in the FigShare repository (https://doi.org/10.6084/m9.figshare.34163682) under the CC BY 4.0 license. The sequencing reads generated in this study have been deposited in NCBI under the BioProject PRJNA1529990 and Sequence Read Archive (SRA) accession numbers SRR40812125–SRR40812128. The 15 high quality cyanobacterial MAGs assembled in this study are available in NCBI with accession numbers of listed in Table S1.

### Authors’ contributions

ES: Data curation; formal analysis; investigation; visualization; writing – original draft; writing – review & editing. BD: Investigation; writing – review & editing; LC: Conceptualization; funding acquisition; supervision; writing – review & editing.

## Supplementary Data Legends

**Table S1**. List of the 269 cyanobacterial taxa included in this study

**Table S2**. ANI and AAI of 15 cyanobacterial MAGs compared with 9,026 published cyanobacterial and melainabacterial genomes

**Table S3**. Genome statistics and completeness of *Gloeobacter dolomitensis* and four published Gloeobacter genomes

**Table S4**. COG annotations of 64 genes specific to the *Gloeobacter dolomitensis* genome

**Table S5**. Structural alignment scores of the SBP located close to the *cynS* of *Gloeobacter dolomitensis* against the reference proteins of various SBPs

**Table S6**. Estimated horizontal gene transfer events among bacterial MAG phyla from Piora Valley dolomite rock samples

**Table S7**. Inferred horizontal gene transfer events, including donor and recipient, among the 15 cyanobacterial MAGs

**Table S8**. Predicted horizontal gene transfer direction among Gloeobacterales genomes, with putative gene names and COG-based functions

**Table S9**. Mean divergence times and 95% HPD intervals of the 15 cyanobacterial MAGs

**Figure S1**. Maximum likelihood phylogeny of the 269 taxa reconstructed from 51,579 unambiguously aligned sites under the best-fit model (LG+F+I+R10) and 1,000 ultrafast bootstrap replicates. Only nodes with bootstrap support values <90% are shown.

**Figure S2**. Genes of interest associated with phycobilisomes, photosynthesis, and respiratory machinery across 18 Gloeobacterales MAGs

**Figure S3**. UpSet plot showing shared and genome-specific orthogroups across five *Gloeobacter* genomes

**Figure S4**. COG functional classification of 64 genes specific to *Gloeobacter dolomitensis*

**Figure S5**. Inferred number and direction of horizontal gene transfers between phyla among MAGs from Rock2-3 dolomite, Piora Valley, Switzerland

**Figure S6**. Inferred number and direction of horizontal gene transfers between phyla among MAGs from Rock4 dolomite, Piora Valley, Switzerland

**Figure S7**. Inferred number and direction of horizontal gene transfers between phyla among MAGs from Rock5 dolomite, Piora Valley, Switzerland

**Figure S8**. Inferred number and direction of horizontal gene transfers between families among cyanobacterial MAGs from dolomite rocks of Piora Valley, Switzerland

