## Supplementary figures and images for "An endolithic *Gloeobacter* and other novel cyanobacterial diversity in alpine dolomites of Piora Valley"

### Figure S1

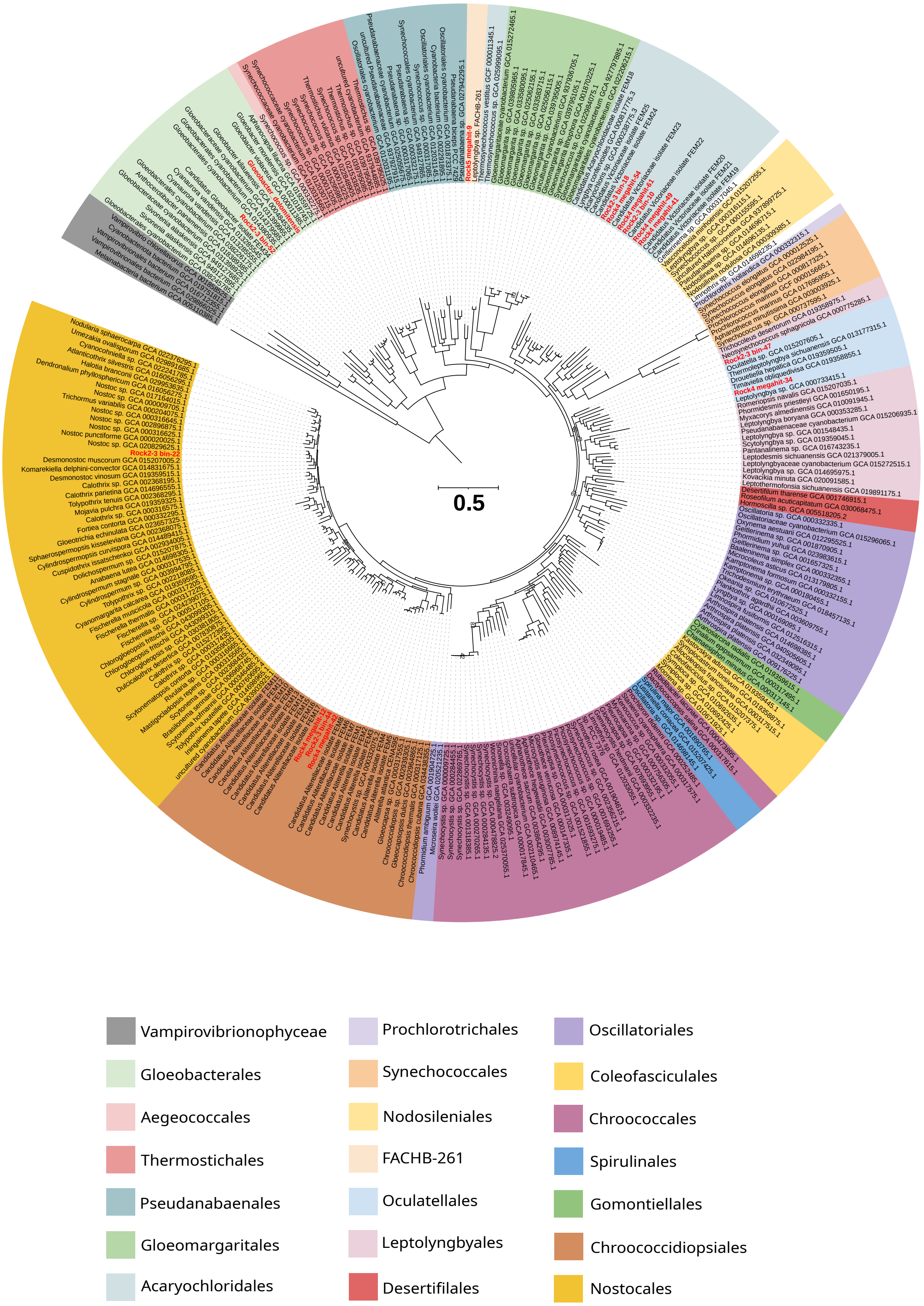

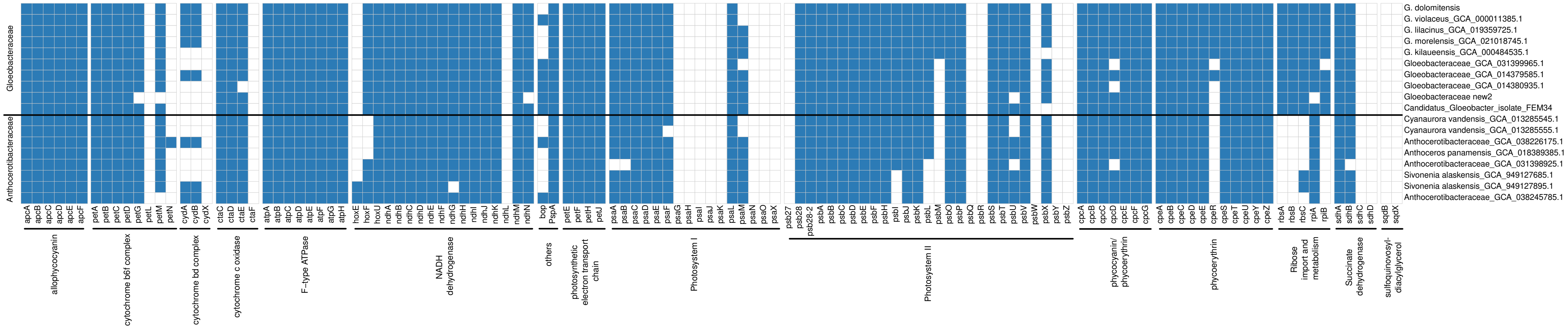

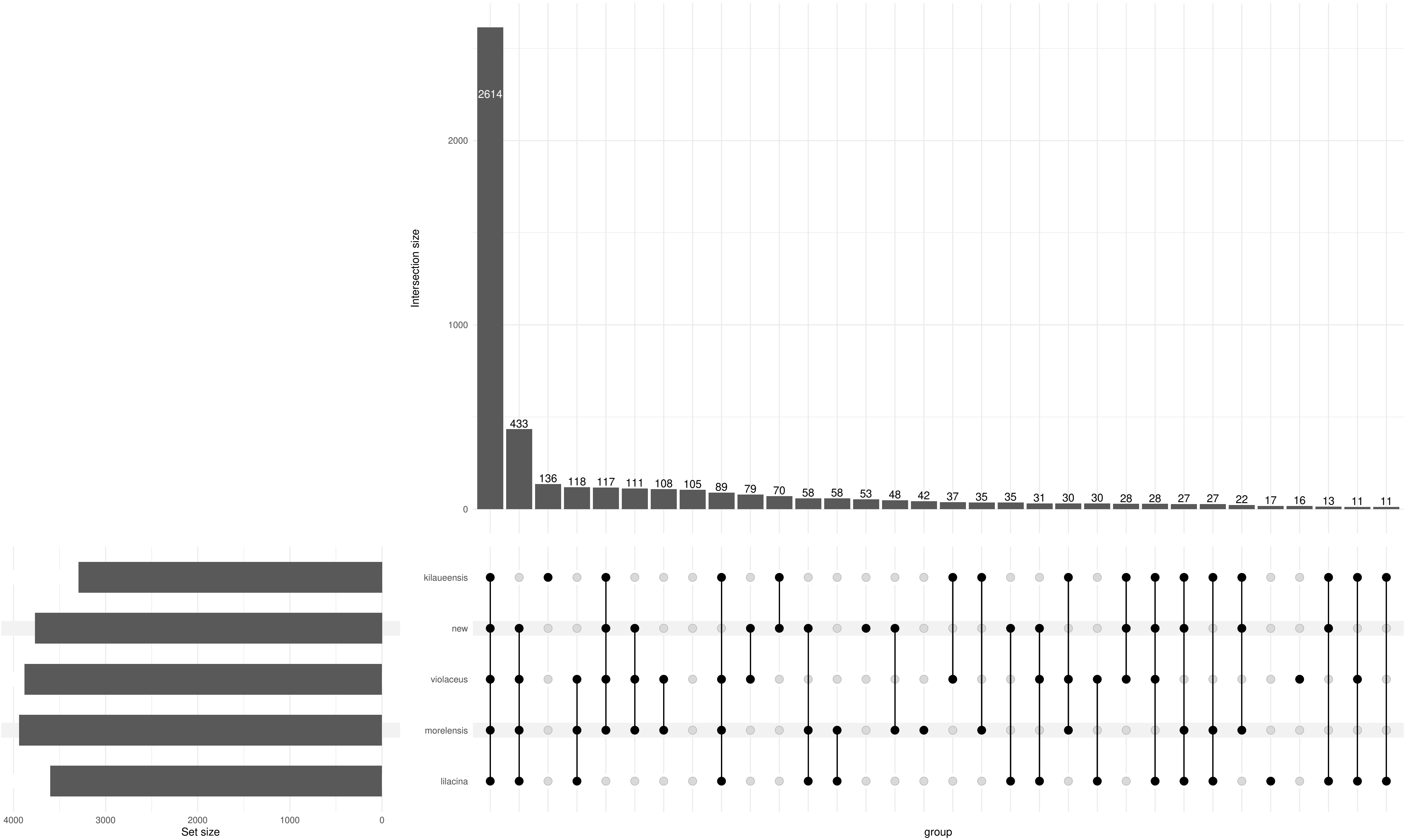

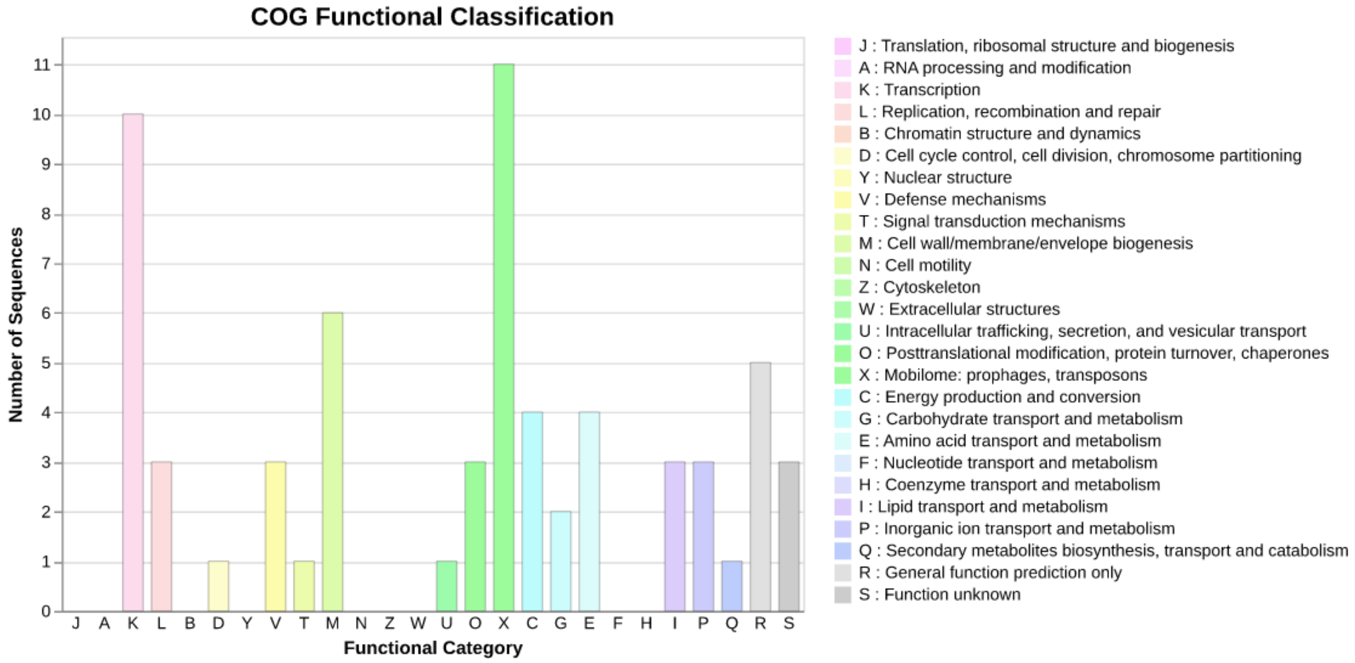

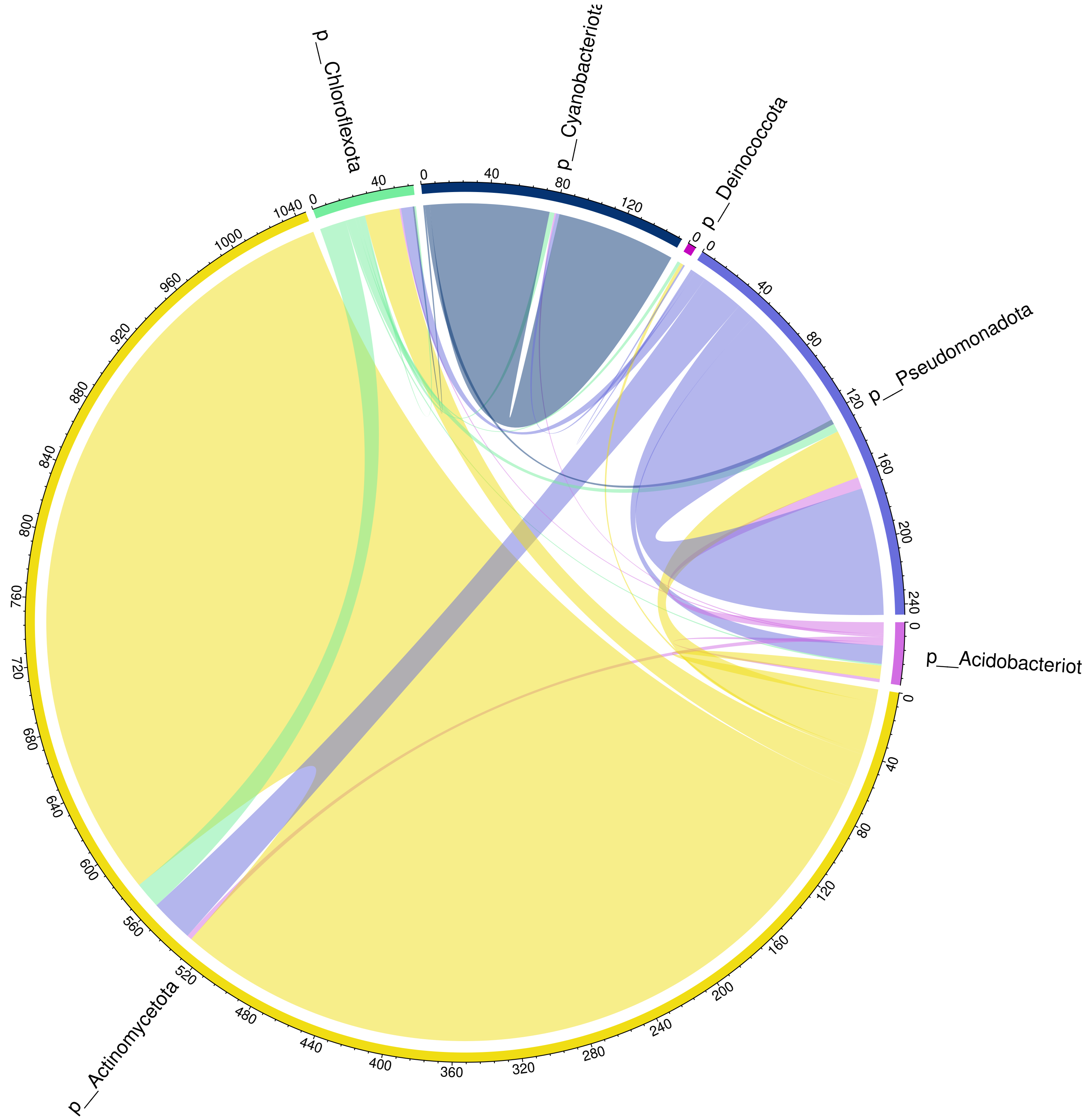

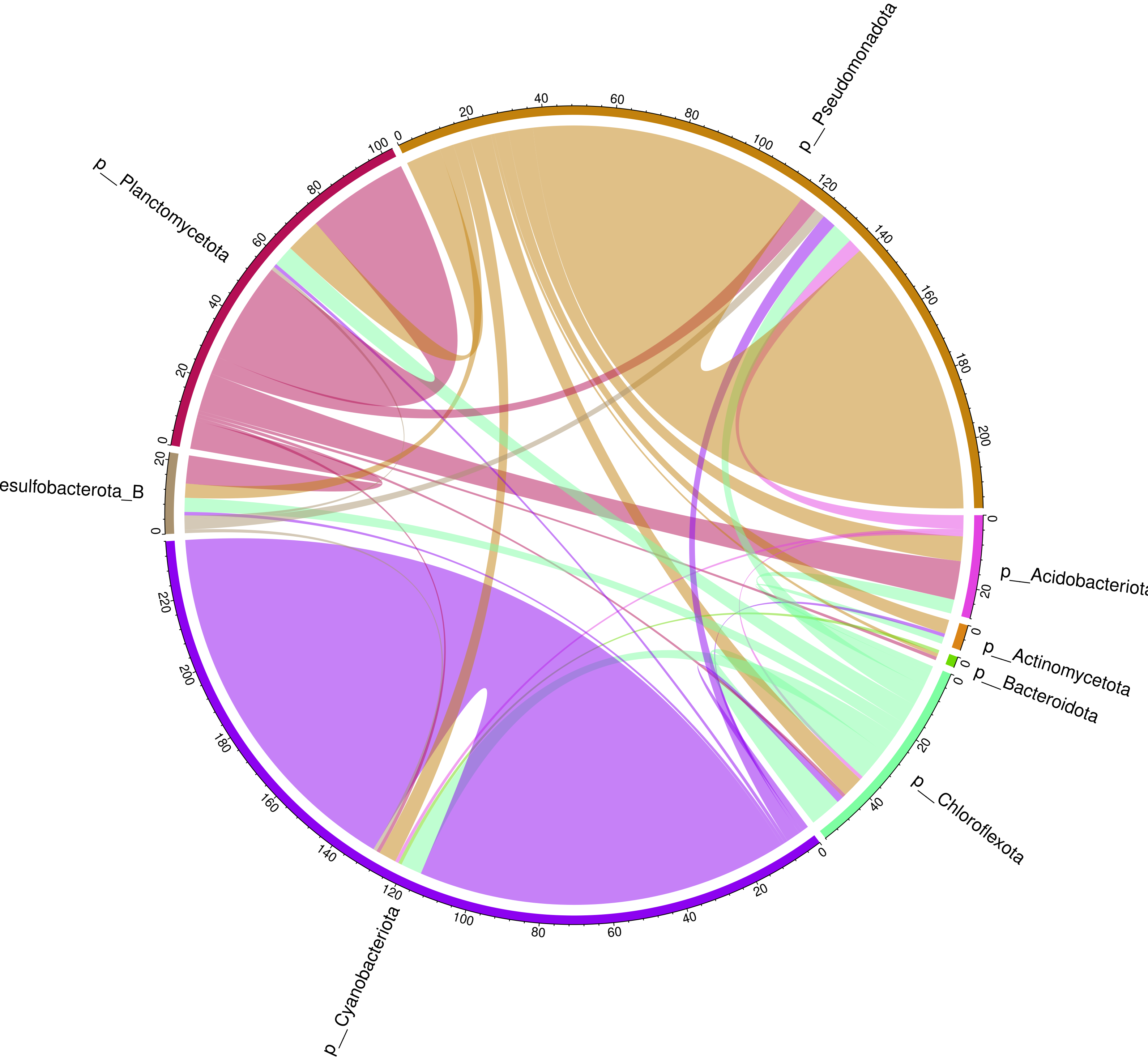

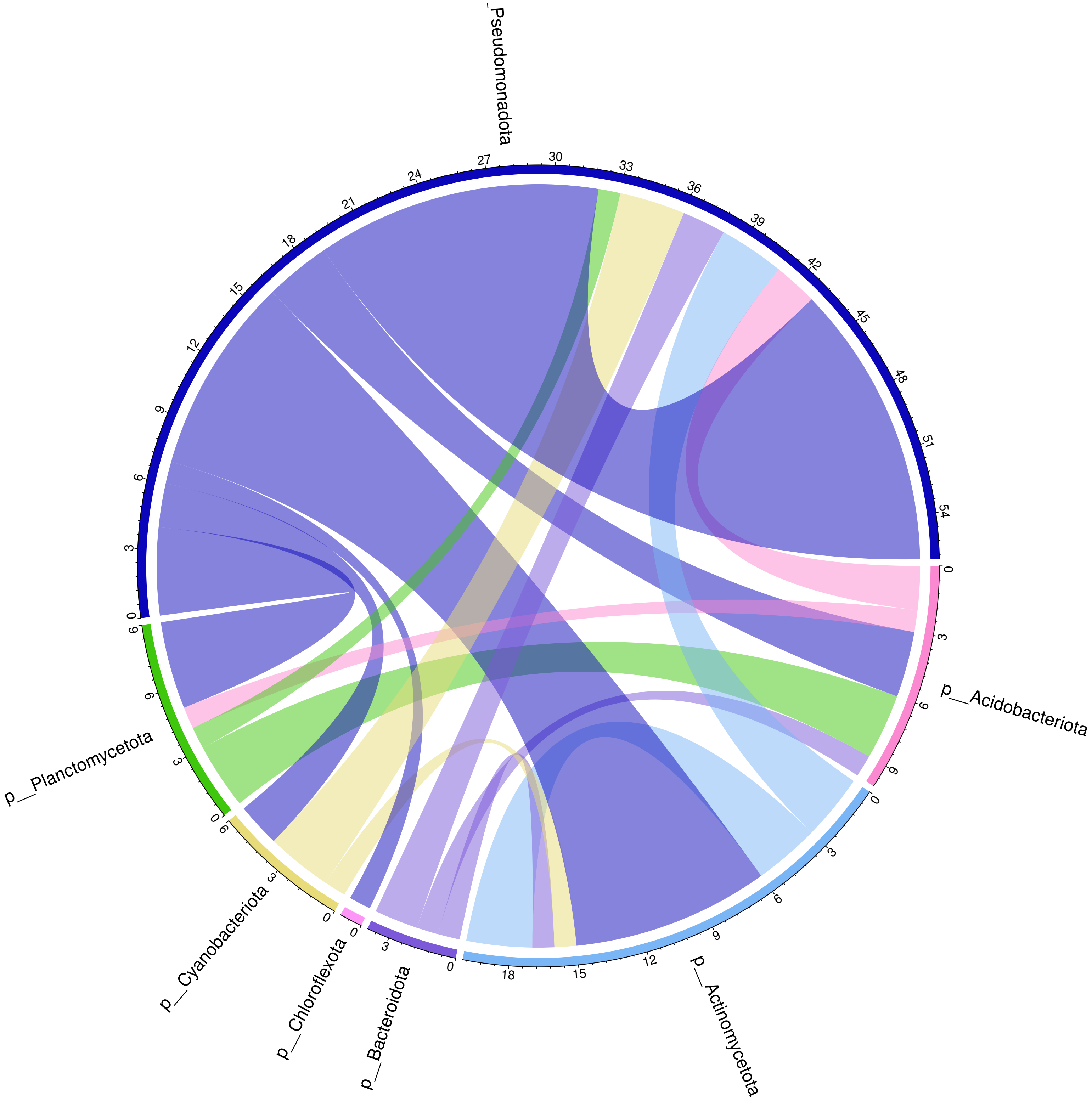

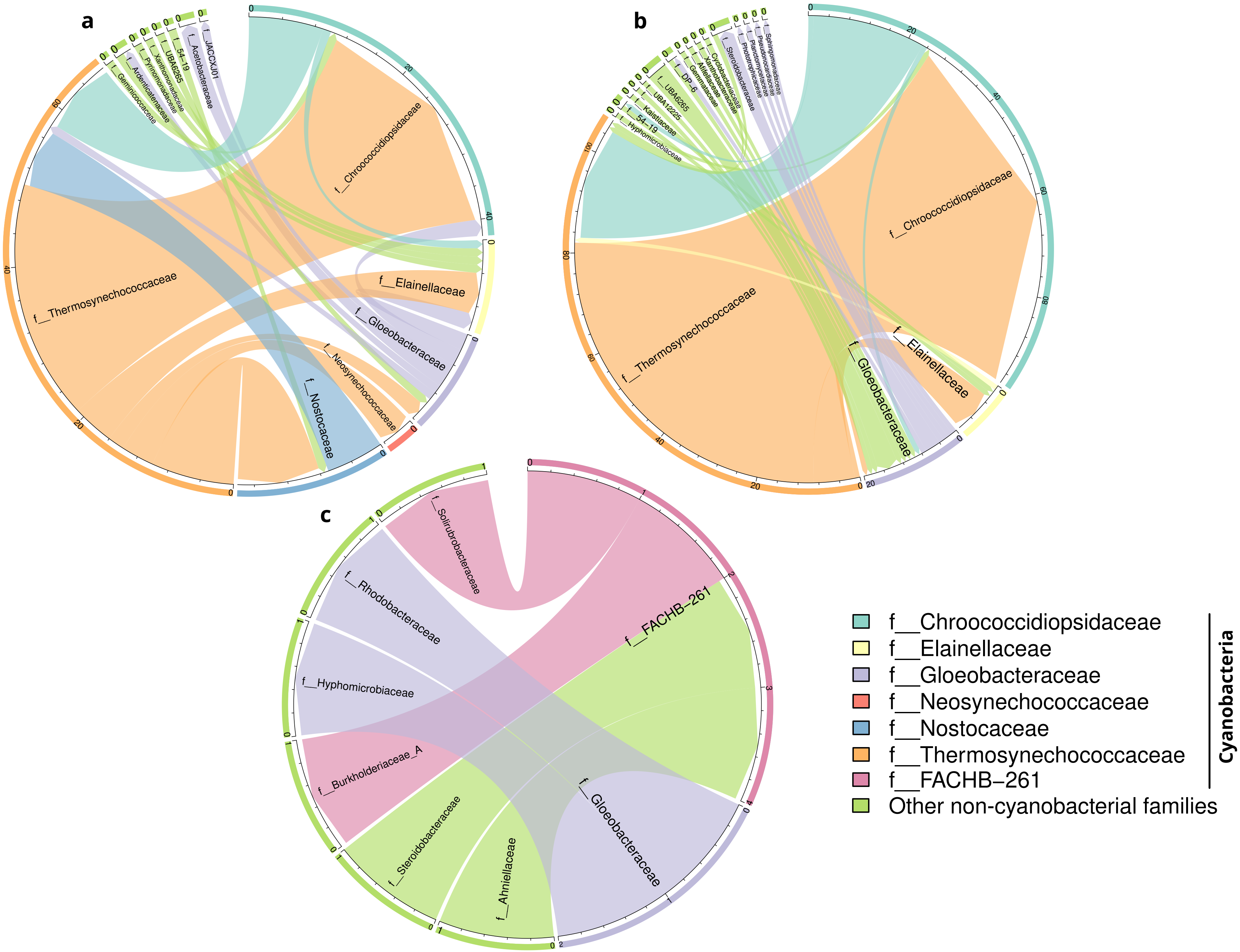
